# Nectar bacteria stimulate pollen germination and bursting to enhance their fitness

**DOI:** 10.1101/2021.01.07.425766

**Authors:** S. M. Christensen, I. Munkres, R. L. Vannette

## Abstract

For many flower visitors, pollen is the primary source of non-carbon nutrition, but pollen has physical defenses that make it difficult for consumers to access nutrients. Nectar-dwelling microbes are nearly ubiquitous among flowers and can reach high densities, despite the fact that floral nectar is nitrogen limited, containing only very low concentrations of non-carbon nutrients. Pollen contains trace micronutrients and high protein content but is protected by a recalcitrant outer shell. Here, we report that a common genus of nectar-dwelling bacteria, *Acinetobacter*, exploits pollen nutrition by inducing pollen germination and bursting. We use time course germination assays to quantify the effect of *Acinetobacter* species on pollen germination and pollen bursting. Inoculation with *Acinetobacter* species resulted in increased germination rates within 15 minutes, and bursting by 45 minutes, as compared to uninoculated pollen. The pollen germination and bursting phenotype is density-dependent, with lower concentrations of A. *pollinis* SCC477 resulting in a longer lag time before the spike in germination, which is then closely followed by a spike in bursting. Lastly, *A. pollinis* grows to nearly twice the density with germinable pollen vs ungerminable pollen, indicating that their ability to induce and exploit germination plays an important role in rapid growth. To our knowledge, this is the first direct test of non-plant biological induction of pollen germination, as well as the first evidence of induced germination as a method of nutrient procurement, as the microbes appear to hijack the pollen’s normally tightly controlled germination mechanisms for their benefit. Our results suggest that further study of microbe-pollen interactions may inform many aspects of pollination ecology, including microbial ecology in flowers, the mechanisms of pollinator nutrient acquisition from pollen, and cues of pollen germination for plant reproduction.

## Introduction

Pollen is an essential food source for many insects, birds, mammals and microbes (1–3), but pollen’s chemical and physical characteristics render it highly resistant to degradation and can make it difficult for consumers of pollen to access the nutrients inside (4,5). As a result, specialized adaptations are required to access nutrients within pollen. However, mechanisms of pollen nutrient acquisition have received limited empirical attention and remain a fundamental enigma in pollination biology, even for the economically important and well-studied pollinators, such as honey bees (6–8).

Pollen is nutrient-dense, but its rich protoplasm interior is enclosed by multiple protective outer layers. The outermost layer, a protective shell called the exine, is made of sporopollenin, a highly crosslinked polymer (9,10) and one of the most highly resistant and enzymatically inert polymers found in nature (4,5). Directly beneath the exine shell is the intine layer, comprised of primarily pectins and cellulose in a multilayered structure; this layer swells and emerges through the exine during germination to form the pollen tube (Fig 1C) (5,11). When pollen germinates (typically on the female stigma), the intine layer is exposed, and is more vulnerable to chemical, osmotic, and enzymatic rupture (Fig 1) (12). The protoplasm is enclosed by the intine, and contains the majority of the pollen nutrition, although the lipid-rich pollenkitt, a waxy layer covering the exine of some pollens, may also contain nutrients (1).

**Fig 1.**
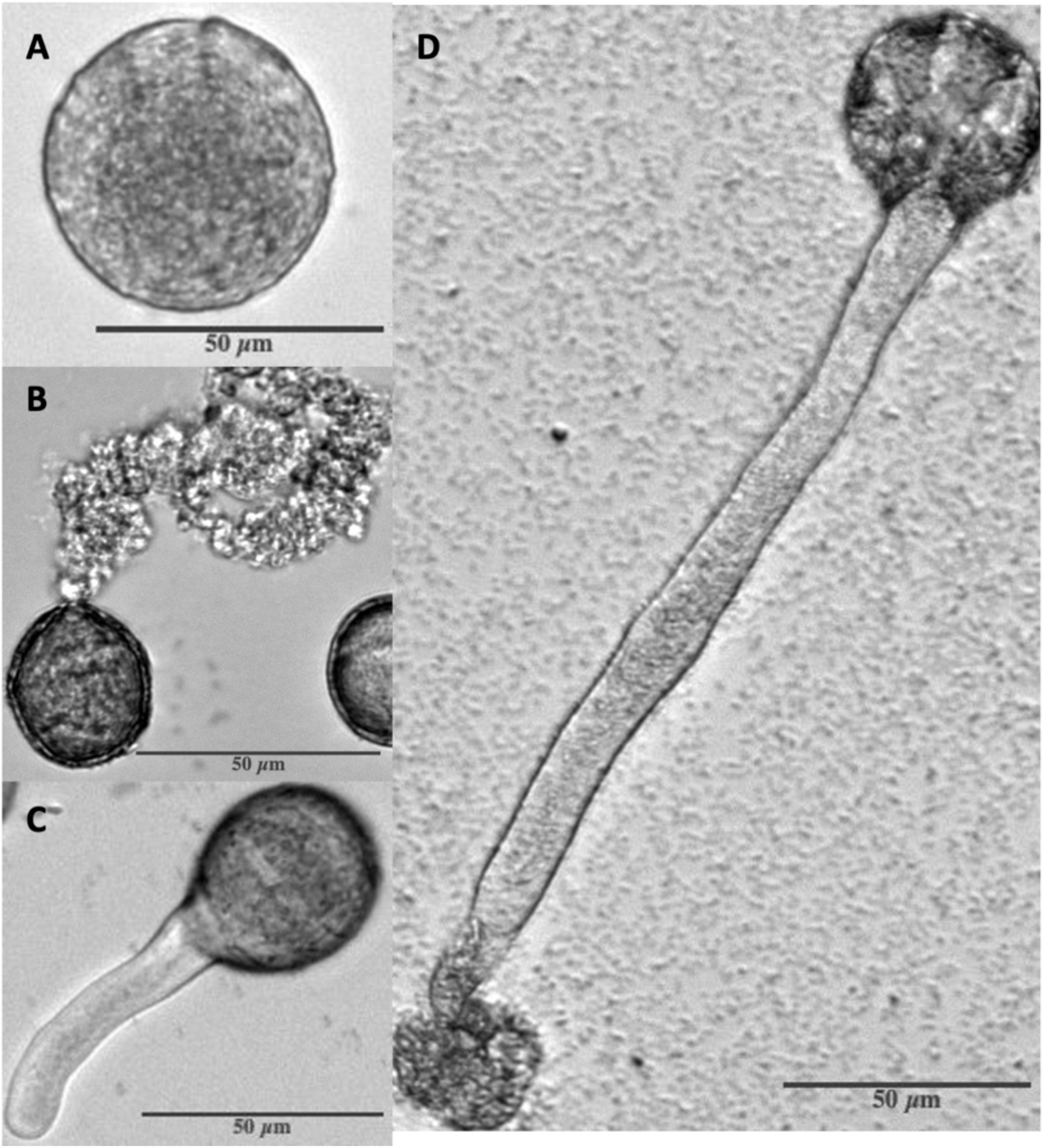
Observed states of *Eschscholzia californica* (California poppy) pollen grains. (A) Whole, ungerminated pollen grain. (B) Burst pollen grain releasing protoplasm without signs of pollen tube. (C) Germinated pollen grain. (D) Pollen grain which has germinated and subsequently the tip of the pollen tube burst, releasing protoplasm.

Although bees are well-known for their consumption of pollen, other organisms, including those found on flowers, also frequently obtain nutrients from pollen. In floral nectar, microbes are thought to be nutrient-limited (14–16), but pollen is commonly found in nectar (17), and can increase microbial population growth rates of nectar microbes (13). Yeasts have been observed aggregating on compromised pollen grains in nectar, suggesting that nectar inhabiting microbes may utilize pollen as a nutrient source (17,18), but mechanisms of nutrient acquisition remain unknown. Nectar-inhabiting gammaproteobacteria in the genus *Acinetobacter* are ubiquitous nectar-specialists (19,20), and known to reach very high (up to 10^9^ cells/ml) densities in nectar despite the nectar’s nutrient imbalance (21–23). Initial observations that *Acinetobacter* grow poorly in traditional media and exhibit specific interactions with pollen grains led us to investigate their effects on one another in more detail.

In this study, we tested the hypothesis that nectar-dwelling and bee-associated *Acinetobacter* species induce pollen germination to access pollen nutrients, and if this phenotype provides them a growth advantage. To test this hypothesis, we examined the interactions between multiple species of the floral and bee-associated *Acinetobacter* and pollen (Table 1). To examine if this ability is shared among common nectar or plant-associated microbes, we also examined *Metchnikowia reukaufii*, a common nectar-inhabiting yeast, and *Pectobacterium carotovorum*, a plant associate and pathogen that produces pectinases. Moreover, pectinases have been implicated in pollen degradation (24,25) and we therefore included *P. carotovorum* because of its high and consistent pectinase production. In addition, we tested whether *Acinetobacter* population growth would benefit from pollen addition, and specifically from germination of the added pollen. Finally, in an effort to explore the mechanism of the observed effects, we assessed pectinase activity to determine if it was associated with microbial effects on pollen. The evidence presented here suggests that nectar-dwelling *Acinetobacter* have a unique ability to induce and benefit from pollen germination and bursting.

**Table 1:**
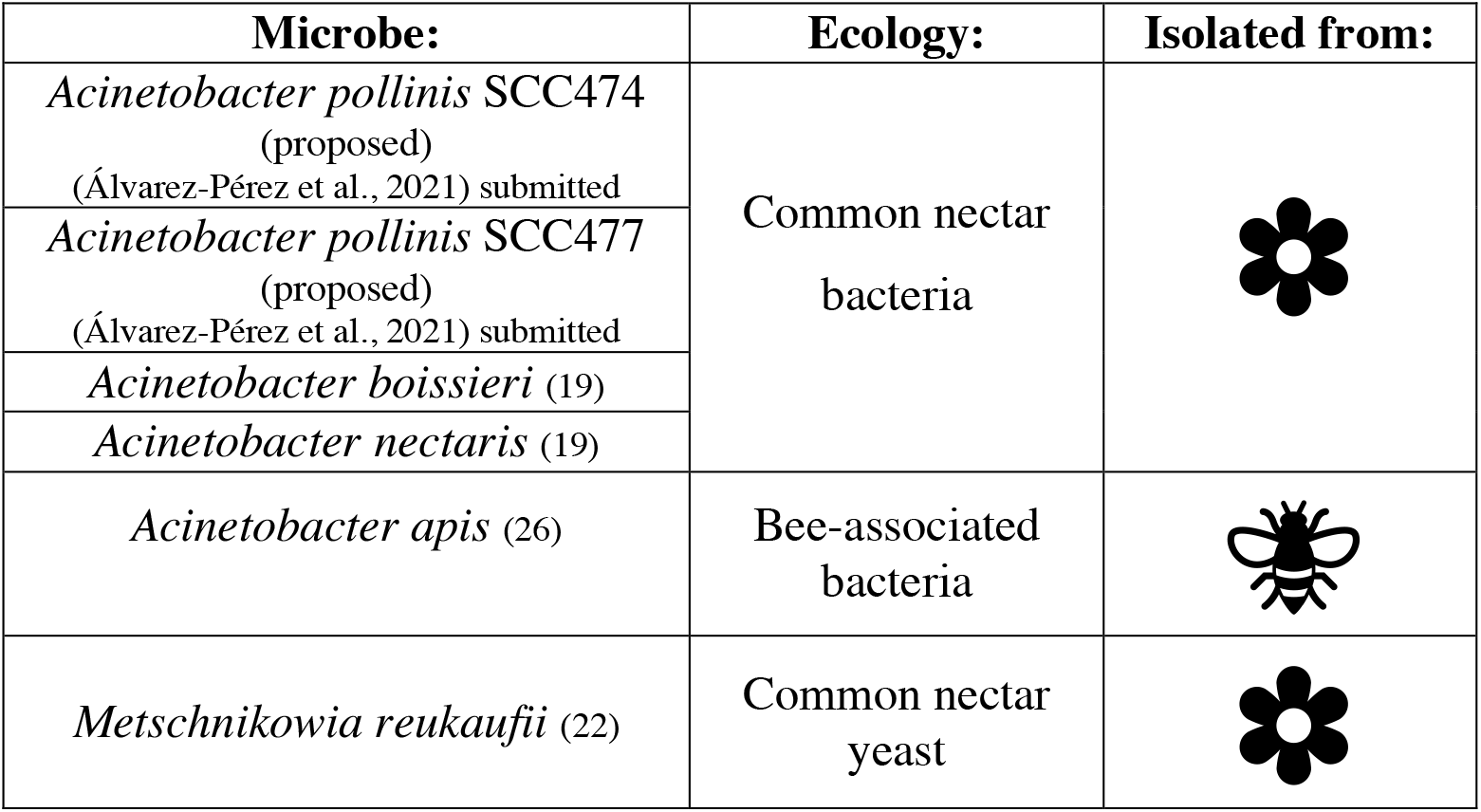
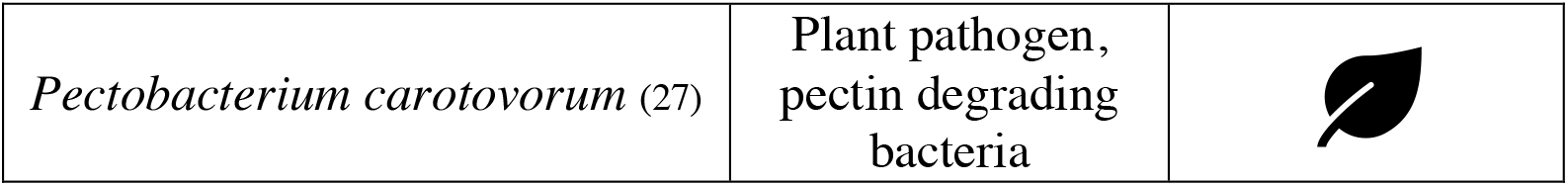
Microbes utilized in germination assays.

| Microbe: | Ecology: | Isolated from: |
| --- | --- | --- |
| *Acinetobacter pollinis* SCC474<br>(proposed)<br>(Álvarez-Pérez et al., 2021) submitted | Common nectar<br>bacteria |  |
| *Acinetobacter pollinis* SCC477<br>(proposed)<br>(Álvarez-Pérez et al., 2021) submitted |  |  |
| *Acinetobacter boissieri* (19) |  |  |
| *Acinetobacter nectaris* (19) |  |  |
| *Acinetobacter apis* (26) | Bee-associated<br>bacteria |  |
| *Metschnikowia reukaufii* (22) | Common nectar<br>yeast |  |
| *Pectobacterium carotovorum* (27) | Plant pathogen,<br>pectin degrading<br>bacteria |  |

**Table 2.** Outline of assays conducted in this study

| Assay name | Strains used | Description | Treatment variable | Response variable |
| --- | --- | --- | --- | --- |
| *Microbial impact (Fig 2)* | All (Table 1) | Visual observation of pollen under microscope over | Microbial species/strain inoculated | Pollen germination |
| *Density dependence*<br>(Fig 3) | *A. pollinis*<br>SCC477 | time after inoculation.<br>Quantification in FIJI. (Supp methods 1) | Concentration (dilution) of *A. pollinis* SCC477 inoculated | and bursting (Fig 1)<br>(percent of total pollen) |
| *Growth benefit*<br>(Fig 4) | *A. pollinis* SCC477, *M. reukaufii* | Direct cell counts (hemocytometer) after 24 hours of growth. | Pollen type/ presence: (none, ungerminable, germinable) | Growth of microbe (cells/ml) |
| *Pectinase assay* (Fig S2) | All (Table 1) except *A. apis* | Plate based, visual qualitative (Y/N) polygalacturonase activity assay. | Microbe species/strain and pollen type/ presence. | Clearance zone on PGA plate (Y/N) |

## Results

### Microbial species and strains differ in effects on *E. californica* pollen germination and bursting

To assess potential effects of microbes on pollen germination, we inoculated pollen solutions and imaged them at 15, 45, and 90 minutes. A grain was counted as germinated if intine was visibly bulging out of the exine, regardless of whether the pollen tube had also burst (see Fig 1 for examples). We observed that by 15 minutes, pollen in wells inoculated with *Acinetobacter pollinis* SCC477 (proposed) or *Acinetobacter boissieri* had on average at least 3 times higher germination than the control uninoculated pollen, pollen inoculated with *P. carotovorum*, or pollen inoculated with *M. reukaufii* (p<0.001 for all comparisons; Fig 2A). Across all treatments, pollen germination increased over time, but wells treated with *A. pollinis* SCC477 and *A. boissieri* had, respectively, 4.4 and 3.7 times higher average pollen germination than the uninoculated pollen at 90 minutes (Fig 2A). In contrast, neither the yeast *M. reukaufii* nor pectinase producer *P. carotovorum* affected pollen germination differently from the uninoculated control at any timepoint. All *Acinetobacter* strains induced significantly higher germination compared to the control by 90 minutes except for *A. nectaris*, which had variable effects on pollen germination (Fig 2A).

**Fig 2.**
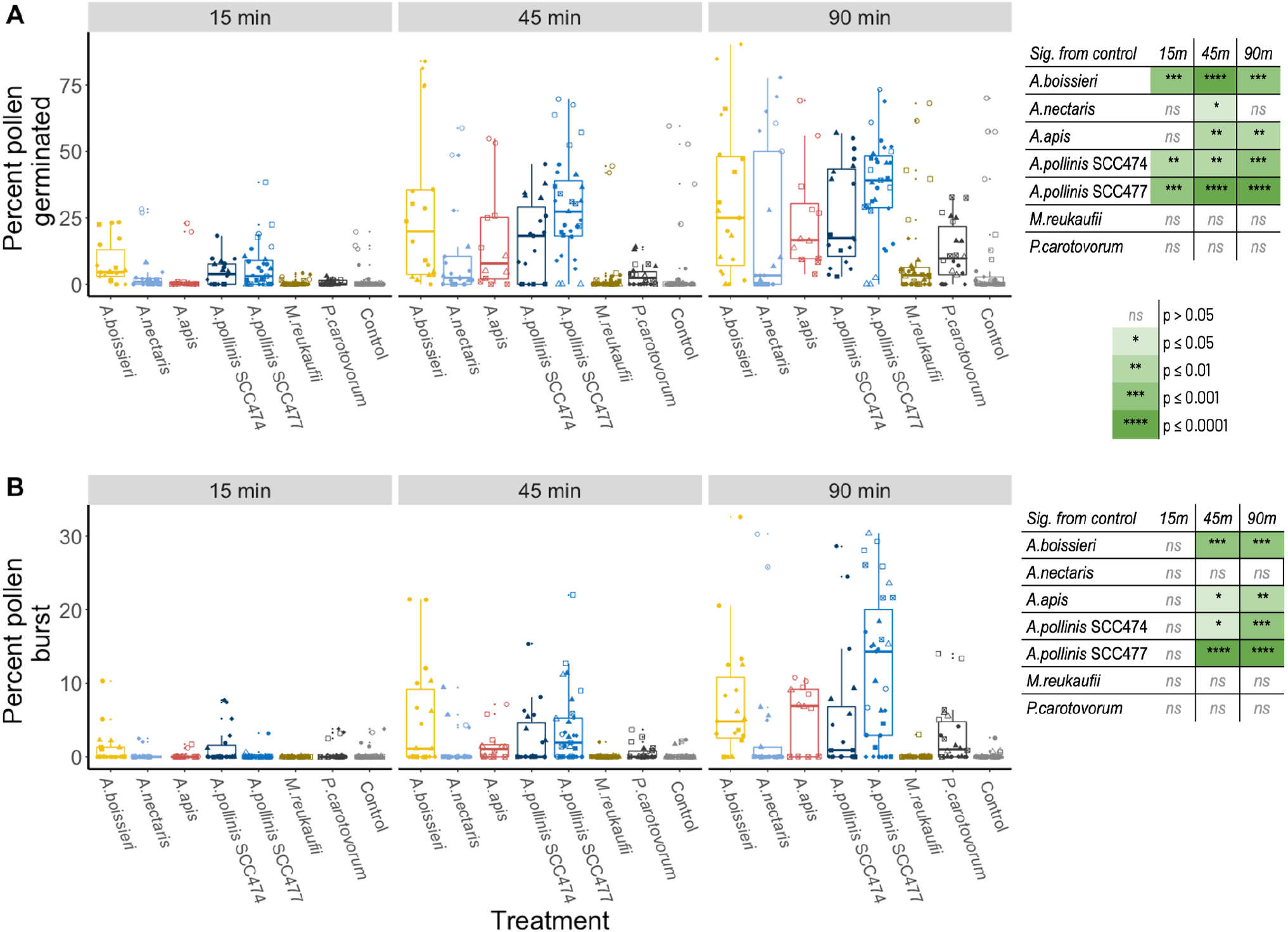
*Acinetobacter pollinis* and *A. boissieri* significantly increase pollen germination and bursting. Proportion germinated includes both intact germinated grains and grains that germinated and subsequently the tip of the pollen tube burst. Proportion burst includes both grains that burst with no visible pollen tube, and grains that germinated and subsequently burst. All data collected at 15, 45 and 90m timepoints-offsets are for clarity. Points represent the mean, with standard error of the mean. Significance from control used Kruskal Wallis test followed by Dunn’s multiple comparisons, with significance cutoff p<0.05. Mean values are based on N=27-36, and a total 41,079 pollen grains.

To assess if and which microbes induce pollen bursting (release of protoplasm, see Fig 1), we also monitored pollen tube and whole pollen bursting visually in the same assays referenced above for pollen germination. A pollen grain was counted as burst if it released protoplasm either from the pollen tube (tip bursting) or directly from the pollen grain (Fig 1B, D). Both *A. pollinis* SCC477 and *A. boissieri* -inoculated pollen showed significantly higher pollen bursting than the no microbe control, *M. reukaufii* (all p<0.001) and *P. carotovorum* (both p<0.05) by 45 minutes (Fig 2B). The average bursting rates for *A. pollinis* SCC477 and *A. boissieri* were 75x and 45x higher than the no-microbe control, respectively. At 90 minutes, pollen inoculated with *A. boissieri, A. apis, A. pollinis* SCC477, and *A. pollinis* SCC474 were burst to a greater extent than in the no microbe control (p<0.001, Fig 2B) but *A. nectaris, A. apis, M. reukaufii*, and *P. carotovorum* did not differ significantly from the no microbe control at any timepoint (Fig 2B).

### Response of pollen to *Acinetobacter pollinis* SCC477 is density dependent

To determine whether there was an essential density of *Acinetobacter* necessary for induction or time until induction of germination or bursting, we conducted a pollen inoculation assay with a range of dilutions of *Acinetobacter pollinis* SCC477 solution. This strain was chosen for further analysis based on high and repeatable difference from the controls in both germination and bursting assays. For both pollen germination and pollen bursting, the response of pollen to solutions of *Acinetobacter pollinis* SCC477 is density dependent. The highest density elicited the most rapid germination (Fig 3A) and bursting response (Fig 3C), and time until germination was longer with lower SCC477 inoculation densities. However, even the lowest density tested elicited germination (Fig 3A) and bursting (Fig 3C) to a similar degree as higher densities (at 24h), but the induction took hours instead of minutes. We calculated ‘T_1/2 max_’-the time it took each treatment to reach half of its maximum germination, for each replicate of the experiment. All treatments except for the uninoculated control reached a max germination of around 40%, varying slightly but not significantly between inoculated treatments. The T_1/2 Max_ increased with decreasing microbial density (Fig 3B, Supplementary code and data); the treatment with the highest density took 40min to reach half of maximum germination, whereas the lowest density took 6hr 13min. The uninoculated control only reached an average max germination of 10% by 24hr, and was quite variable between replicate wells, with two of the three wells barely germinating at all.

**Fig 3.**
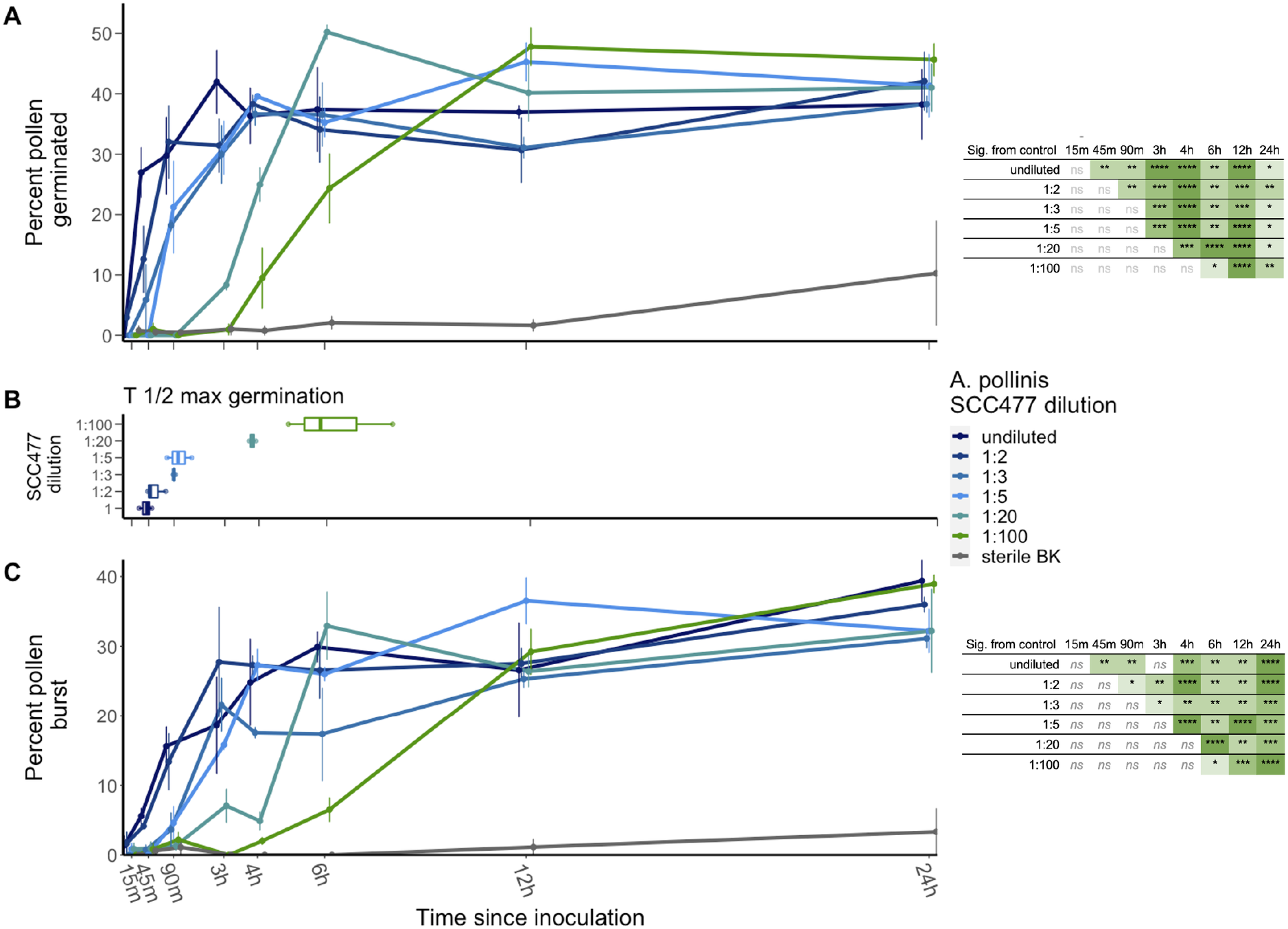
Response of pollen to *Acinetobacter pollinis* SCC477 is density dependent. Points represent the mean of 3 biological replicates of each density of *A. pollinis* SCC477, and a total of 7,618 pollen grains, an average of 952 per timepoint. Standard error bars are shown. A) Germination over time for pollen inoculated with dilutions of SCC477 from 1-1:100, boxplot indicates time to half-maximum germination (∼40%) for inoculated wells. B) T_½ max_ for each density. C) Bursting over time when inoculated with dilutions of SCC477. Undiluted indicates a OD600=0.1 solution. All data collected at labelled timepoints-offsets are for clarity. Significance from control (sterile BK) calculated with ANOVA followed by Tukey multiple comparisons.

Similarly, pollen burst at earlier timepoints with high microbial density than the lower densities (Fig 3C). For all treatments, bursting lags behind the respective germination times (Fig 3C). In addition, pollen tube length, an indication of time elapsed between germination and bursting or halting of growth also responded to microbial density, with longer tubes observed in the lowest-density treatments (1:20, 1:100) compared to shorter tubes in the most concentrated (1, 1:2) (Fig S1).

### *Acinetobacter pollinis* SCC477, but not *M. reukaufii*, benefits from germinability of pollen

To assess whether pollen presence or germination affects microbial growth, we assessed the growth of two strains that differed in their ability to germinate pollen--*A. pollinis* SCC 477, which readily germinates pollen and the yeast *M. reukaufii* which is unable to germinate pollen. To microbial suspensions, we added either no pollen, germinable pollen (GP,) or pollen rendered unable to germinate (ungerminable, UP). Compared to their growth with no pollen, *A. pollinis* SCC477 and *M. reukaufii* displayed significantly higher cell density after 24h with the addition of any pollen (Fig 4). However, only *A. pollinis* SCC477 growth differed with pollen germinability: its cell density after 24h was almost twice as high with germinable pollen (1.9x higher than UP, 7.4x higher than no pollen control, avg 4.6×10^8^ cells/ml) than it was with ungerminable pollen (3.8x higher than no pollen control, avg 2.3×10^8^ cells/ml) (p<.0001) (Fig 4A). Moreover, in the absence of pollen, *A. pollinis* SCC477 barely grew from the starting concentration (∼5×10^7^ cells/ml) during the 24h window (avg 6.2×10^7^ cells/ml) (Fig 4A), indicating significant reliance on pollen-derived nutrients. *M. reukaufii* grew equally well with both pollen treatments. Upon visually checking the pollen grains at 24h, we confirmed that our treatment was effective in preventing the pollen’s germination in the “ungerminable” treatments.

**Fig 4.**
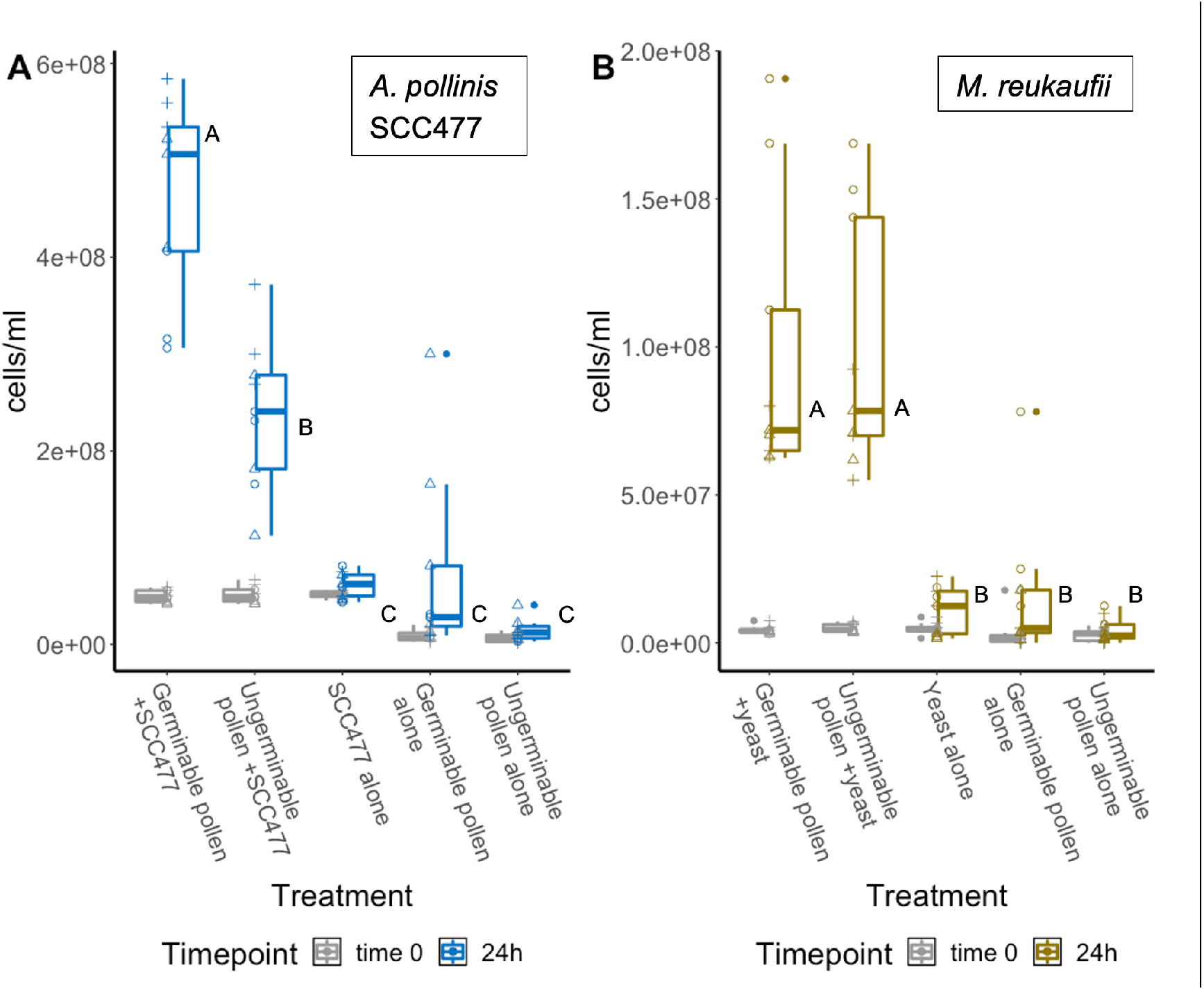
*Acinetobacter* growth is enhanced by germinable pollen addition. Standard error bars are shown, significance between treatments marked by lettering. Data point shape indicates assay replicate. A) *A. pollinis* SCC477 data represents three assays, 3 biological replicates of each treatment per assay, N=9. B) *M. reukaufii* data represents three assays, 3 biological replicates of each treatment per assay, N=9.

### Pectinase activity is not associated with microbial effects on pollen

Pectinase activity is inferred to play a role in pollen degradation. To determine if pectinase activity varied among microbial species and strains, we used a plate-based polygalacturonase (PGA) assay to detect pectin degradation, as polygalacturonic acid is the backbone of pectins. We also tested whether pollen exposure activated pectinase activity. While the known pectinase producer, *P. carotovorum* was able to consistently and clearly degrade PGA, *Acinetobacter* strains were either unable to do so or showed very faint degradation halos in only a few replicates (Table S1). Further, *A. pollinis* SCC477 did not show clear or repeatable clearance zones even after prior exposure (1h) to pollen (Table S1, Fig S2).

## Discussion

The mechanism of pollen digestion and nutrient acquisition remains a fundamental enigma in pollination biology. Five general mechanisms have been suggested: mechanical disruption, piercing, enzymatic or chemical penetration, osmotic shock, and induction of germination (1). Here, we show evidence that a genus of common nectar- and bee-associated bacteria can induce pollen germination, compromise pollen’s protective outer layers, and gain access to the protoplasm. Although this mechanism had been previously suggested as a potential method in macroscopic pollen consumers (8,28,29), direct induction of germination has not been empirically documented. Though access to protoplasm via pollen bursting is the most apparent benefit of induced germination, the process of germination can stimulate pollen to release free amino acids and proteins (30), providing nutrients even without direct protoplasm access.

Pollen germination can be highly specific, being influenced by hydration rate of the desiccated grain (31), pectin modifications (32), pistil-pollen crosstalk (33), calcium (34), and other molecules (35). It is generally observed that pollen germination *in vitro* is much slower than stigmatic germination, presumably due to missing factors or less-than-ideal conditions (33). The mechanism behind the induced germination phenotype described here has yet to be determined but dissecting this novel phenotype may be useful in developing further understanding of the mechanisms that promote and regulate pollen germination and bursting, especially if *Acinetobacter-*induced germination and bursting is shown with pollen of additional plant species. Moreover, the variability within and among strains of *Acinetobacter* (Fig 2) could reflect species specificity in interaction or reliance on pollen (36) (eg. *A*.*pollinis* SCC477 interacts with *Eschscholzia* pollen, whereas other *Acinetobacter* species may interact more strongly with other pollen genera), recent evolution of the phenotype, or uncharacterized environmental pollen variation (eg. humidity, temperature, low abundance microbes or endophytes). Further testing on other pollens under various conditions, as well as characterization of the mechanism, will be necessary to determine the biological underpinnings of the observed variation.

We had hypothesized that pectinase activity would underlie this phenotype, as pectinases have been implicated in facilitation of pollen digestion in other studies (24,37,38), and purified pectinase enzymes are known to induce pollen germination at low densities and bursting at higher concentrations (39). Annotated genomes of nectar *Acinetobacter* have multiple genes encoding proteins assigned as pectinases in their genomes (accession numbers and locus tags in Table S1).

However, two lines of evidence suggest that pectinase activity may not play as strong of a role as predicted. First, the known pectinase producer *Pectobacterium carotovorum* did not significantly affect pollen germination or bursting despite observed pectinase activity in the PGA assay. Second, we detected no or minimal levels of PGA degradation by *Acinetobacter* strains, even when exposed to pollen. This is notable because it is commonly assumed that pectin degradation genes are associated with pollen digestion (24,37). Although pectinases may be involved in some stages of pollen degradation, our data suggest that pectinases may not be solely responsible for this phenotype. Alternatively, different pectinases may exhibit varying levels of substrate specificity, or have different requirements for expression; additional gene expression data and enzyme characterization of *Acinetobacter* pectinases would be necessary to test this hypothesis.

The nectar environment is a highly specialized niche, requiring microbial inhabitants to tolerate high osmolarity, plant secondary metabolites or toxins, ROS, and low nutrient levels-all within an ephemeral habitat (40–42). In this environment, the development of mechanisms to access protected nutrients could be essential for the rapid growth needed for further dissemination (43). The ability to induce pollen germination and bursting could help to explain the prevalence of *Acinetobacter* species in floral nectars and other environments frequently containing pollen, as well as the high densities observed by (21–23). This is supported by our *Acinetobacter pollinis* SCC477 growth results, as *A. pollinis* SCC477 grew to nearly double the cell density (1.8x) when given fresh, germinable pollen versus pollen that was unable to germinate over 24 hours. In contrast, *M. reukaufii*, a common nectar yeast, did not induce pollen germination or bursting and its growth did not differ between fresh and ungerminable pollen, presumably because it did not gain access to the protoplasm although it likely accesses nutrients from pollenkitt or leached nutrients. The fact that *M. reukaufii* grew equally well on both pollen treatments also indicates that our treatment to render the pollen ungerminable did not decrease its overall nutritional quality. The difference between microbial species’ effects on pollen may allow for niche partitioning or bacterial effects may facilitate yeast growth (15,36).

There are many ecological implications of microbial effects on pollen that remain to be explored. Since many pollen consumers also consume floral nectar, *Acinetobacter-*induced pollen germination and bursting could illuminate a previously undocumented mechanism for macroscopic species to access pollen nutrients via microbial partners. While pollen germination has been considered and studied as a potential mechanism of digestion for macroscopic consumers(8,28,29), microbial involvement has not been studied. *Acinetobacter spp*. are common in floral nectar, and are also found in the nectar and pollen provisions that bees provide their offspring (44–46). Recent work found that blueberry bee (*Osmia ribifloris)* larvae that receive provisions without this original set of microbes grow more slowly and have lower survival than larvae with the live microbes in their provisions (47). It is also known that pollen grains with protruding intine, as occurs in germination, are more completely digested by solitary bee larvae (48,49) and adult honey bees (50). In a study of pollen modification before and during digestion by the European orchard bee (*Osmia cornuta)*, Nepi et al. found that pollen in the provision had protrusions from the apertures, which they report resembles germinating pollen, without formation of the full pollen tube (49). They conclude that this germination or pseudo-germination could act as a “pre-treatment” for digestion and speculate that commensal microbes may also play a role (49). Our study suggests that the role of floral microbes in pollen germination and bee nutrition warrants further study.

Microbial stimulation of pollen germination may also affect plant fitness. Previous work has shown that nectar microbes can alter pollinator preference for flowers by altering nectar composition and volatiles, directly impacting reproduction (18,51)—effects on pollen nutrient availability or scent could also possibly influence pollinator foraging and pollination, thereby affecting plant reproduction. More direct effects are also possible. *Acinetobacter* is common on many floral surfaces, including stigmas (52) and is prevalent in seed microbiomes (53), particularly following pollinator visitation. Whether *Acinetobacter* growth on stigmas affects pollen germination and success in fertilization will also require further study.

As we have demonstrated, the presence and growth of floral *Acinetobacter* can significantly affect pollen physiology and may have important consequences for plants and pollinators (18,42,47,51). Using members of the nectar-dwelling *Acinetobacter* clade, we have shown that common nectar bacteria can increase the speed and overall proportion of germination and bursting of pollen grains, that these responses are bacterially density-dependent, and that inducing pollen germination benefits bacterial growth. The results presented here suggest that further study of microbe-pollen interactions could shed light on the specific biochemical triggers of pollen germination and plant reproductive success and the ecological effects of microbe-pollen interactions on plants and pollinators.

## Materials and methods

### Study system

We utilized pollen from the California native poppy *Eschscholzia californica* (Papaveraceae) as it produces a large amount of pollen, is easily obtainable near our laboratory to allow for minimal effects of storage, and because flowers are covered by a cap that is pushed off just before its first opening with the sun, so flowers can be collected before insect or wind activity introduces contaminating microbes. Pollen of Papaveraceae has ectoapertures (54,55) meaning that the outermost layer of the exine shell is missing from the aperture region but there are still other sporopollenin layers surrounding the intine (56).

We chose to examine the nectar *Acinetobacter* clade and their impact on pollen after initial observations of *Acinetobacter* –induced pollen germination. We chose 3 nectar-specialist *Acinetobacter* species representing the species most commonly isolated from nectar *(A. boissieri* and *A. nectaris)*, and an additional species in the clade that was isolated from the honeybee gut (*A. apis*). We recently isolated and described *A. pollinis* (Alvarez-Pérez et al. submitted), of which we used two strains. *Metschnikowia reukaufii* is a commonly isolated nectar-dwelling yeast and has been observed to negatively impact germination of *Asclepias* pollinia and aggregate around the pollen of *Helleborus foetidus* (17,57). We included the pectin degrader and plant pathogen *Pectobacterium carotovorum* as a control for high pectinase activity and because it is a plant associate that is not associated with nectar or pollen (27).

### Do Acinetobacter spp and other nectar and plant associated microbes differ in their ability to induce pollen germination and bursting?

To test whether microbial inoculation impacts pollen germination and bursting, we collected newly opened *Eschscholzia californica* flowers the morning of the assay to minimize the chances that flowers had been visited by insects or otherwise “contaminated” with microorganisms or other pollen grains. Many attempts were made to sterilize the pollen prior to the assays (low heat sterilization, autoclaving, chlorine gas fumigation, 70%-50% ethanol submersion, 10%-1% bleach submersion), but all rendered the pollen unable to form visible pollen tubes at all or were not effective at sterilizing pollen, so newly collected pollen from pre-collected flowers was used. Newly collected pollen had low levels of microbial growth (Fig 4).

For all microbial growth assays, microbial strains were plated from freezer stocks onto Tryptic Soy Agar (+5% fructose), then colonies were inoculated into Tryptic Soy Broth and incubated at 30C for 1-3 days. Suspensions were normalized to an OD600 of 0.1 or 0.05 (depending on the microbe with the lowest OD, and consistent within each assay). Suspensions were centrifuged for 5 minutes at 12,000 rpm to pellet the cells, and remove the growth media and any released metabolites and proteins, and cells resuspended in Brewbaker and Kwack (BK) pollen germination media (10% sucrose, 100mg/L boric acid, 300 mg/L calcium nitrate, 200mg/L magnesium sulfate, 100 mg/L potassium nitrate (34)). After resuspension by repeated pipetting, the suspension was vortexed for 30s and then sonicated for 3m. To ensure no carryover of metabolites from microbial growth media, these washing steps (from centrifugation through sonication) were performed twice for the first five assays, and three times for the last four assays. 50ul of each washed microbial BK suspension (or control BK) was added to respective wells (3-5 replicate wells per microbe per assay) in a sterile, flat bottom, clear 96 well plate.

Immediately before beginning each assay, the pollen solution was prepared. Briefly, pollen from 1-3 flowers was vacuumed or shaken directly from anthers into a sterile 5mL Eppendorf tube and 2mL of BK solution was added. Immediately after pollen grains were in solution, 50ul of the pollen solution was added to each well of the sterile 96-well plate, mixing pollen stock solution every 5-7 wells to ensure even pollen deposition in each well. Control (uninoculated) wells also received 50ul of pollen, but instead of 50ul of microbe solution, they received 50ul of sterile BK. The plate was covered with a lid and wrapped in a layer of aluminum foil to limit evaporation, contamination, and light exposure. Each treatment was replicated in at least three wells per assay. Each well of the plate was then imaged on an EVOS M5000 imaging system at 15m, 45m, and 90m after mixing.

### Image analysis

Images were analyzed in FIJI2 (v2.0.0) (58), utilizing a custom Macro script to count total pollen grains in each image (Supplementary methods 1). The numbers of germinated, burst, and tip burst grains were then counted by hand in FIJI2 using CellCounter (59). A grain was counted as “germinated” if the pollen intine was clearly emerging from the exine (Fig 1C), as “burst” if the pollen grain was surrounded in loose cytoplasm, but without any visible protrusions (Fig 1B, Supplemental video), and as “tip burst” if the pollen grain had germinated, and then the pollen tube had burst and released cytoplasm (Fig 1D, Supplemental video). For all assays, pollen grains were classified twice to confirm accuracy; once just after assay completion, and all assays were recounted by the same observer within a week prior to combined final analysis to ensure consistency of each category between assays, as well as validation of previous data. Eleven total assays were done, and two were eliminated from further analysis. One assay was eliminated due to data loss of images-making recount and validation of pollen ratios impossible, and the other was eliminated due to contamination of the initial growth media. An average of 4,564 (+-1501.6) pollen grains were counted per assay, with a total of 41,079 pollen grains overall.

### Germination data processing

For analysis, the pollen grain counts in each category (described above in main assay description) were converted into percentages of the total number of pollen grains in each image. “Germinated” and “tip burst” were combined in the category of “all germination”, because the “tip burst” pollen had also germinated. Likewise, “burst” and “tip burst” were combined to form the category “all burst” because the pollen grains in both initial categories expelled their protoplasm into the solution.

### Is pollen germination/bursting Acinetobacter pollinis SCC477 density-dependent?

To test whether pollen responds differently to differing amounts of *Acinetobacter*, we repeated the above methods, but inoculated pollen with dilution series of *A. pollinis* SCC477 suspensions (1, 1:2, 1:3, 1:5, 1:20, 1:100) as well as sterile BK (no microbes). We started with a solution at an OD of 0.1 (10^7^ cells/ml) and diluted to the above ratios, with the first being “full concentration” and the last consisting of only sterile BK. We then imaged the wells at 15m, 45m, 90m, 3h, 4h, 6h, 12h, and 24h. The images were analyzed, and pollen grains counted and categorized as described for the previous assay, but in addition we measured and recorded the lengths of each pollen tube in all of the images using FIJI2 (Supplementary methods 1).

### Does Acinetobacter pollinis SCC477 benefit from pollen addition and pollen germination?

To test whether *Acinetobacter* benefits from pollen addition and specifically from pollen germination, we monitored growth responses of *A. pollinis* SCC477 in the presence of live pollen, pollen rendered unable to germinate, and without pollen. We created washed suspensions of OD600 0.1 *A. pollinis* SCC477 as above (pellet cells, remove media, replace with BK, vortex, sonicate-repeated 3x). *A. pollinis* SCC477 was selected because its addition resulted in the greatest induction of germination and bursting of the species and strains that we tested. The initial cell number of *A. pollinis* SCC477 cell suspension was immediately counted on a hemocytometer and adjusted to 5×10^7^ cells/mL.

Pollen was collected as above and was weighed out into each of two 5ml tubes. The first was microwaved for 3 minutes to prevent germination by submerging the pollen in BK media, opening the lid and placing in a glass beaker to hold tube upright, then immediately microwaving on high for 3 minutes. The second tube remained untreated. Just before combining at time 0, BK solution was added to the fresh, germinable tube of pollen. All assays had consistent concentrations of pollen (4.2mg/mL).

In 1.7ml Eppendorf tubes, 50ul of *A. pollinis* SCC477 solution was mixed with 50ul of the microwaved pollen solution, the live (untreated) pollen, or the no-pollen (BK) control, and 400 ul of sterile BK (N=3 for each pollen type). We added the same amount (50ul) sterile BK instead of *A. pollinis* SCC477 to 3 tubes each of both untreated and ungerminable (microwaved) pollen as a control for the microbes already present on the pollen. The initial concentration of microbes in each tube was determined by hemocytometer cell count. The tubes were sealed with headspace and left at room temperature for 24h. At 24h each tube was sonicated to separate cells for visual enumeration, as nectar-dwelling *Acinetobacter* can form dense biofilms, then vortexed, diluted 1:10 in PBS, and counted on the hemocytometer. The pollen grains in the tubes that pollen was added to were checked to ensure that the microwaved pollen did not germinate, and that the fresh pollen could. 100ul of each tube was also plated at 24h to verify that bacterial colony morphology and characteristics matched the inoculum. Individual colonies from these plates were correctly identified by MALDI-TOF and Compass Explorer as described below.

To assess if pollen germination affects the growth of microbes that do not induce germination, we repeated this assay with *M. reukaufii*, a common nectar yeast. Instead of beginning with 5×10^7^ cells/mL, we began with 5×10^6^ cells/mL, to control for microbial biomass.

### Microbial identification/validation

Because effective sterilization of the pollen could not be done without compromising germinability, we wanted to confirm that added microbes remained as the primary species/strain in their wells, and to identify pollen-borne microbes in the control wells. We therefore validated microbial identity with MALDI-TOF for some of the assays (Supplementary methods 2)(60). After imaging the wells, each well was plated out onto Tryptic Soy Agar, colonies were isolated and spectra were generated using MALDI-TOF and associated software (Bruker-Ultraflex III, Bruker Daltonic-flexControl v. 3.4) and subsequent spectral analysis and matching in MALDI Biotyper Compass Explorer software (Bruker Daltonic, v. 4.1.70). The spectra were compared to both the pre-loaded Bruker microbial isolate library and our additional custom library of floral microbes (Supplementary data-MALDI_identifications.csv).

### Are pectinases involved in the stimulation of pollen germination?

Microbial strains described above were grown on either R2A or YM plates. For PGA degradation experiments, microbial strains were spotted onto PGA plates containing 0.5% (w/v) PGA (Sigma) in 0.1 M Tris-HCl pH 8.6, with 1.5% agar and 0.5% yeast extract. Plates were incubated for 48-72h at 37°C and stained with 1% (wt/vol) cetyltrimethylammonium bromide (CTAB-Chem-Impex cat. no 01781) for 20 minutes to visualize clearance zones. Each microbe spot was replicated 9 times (Table S1).

To determine whether exposure to pollen would increase or trigger polygalacturonase activity, we grew *A. pollinis* SCC477, *Pectobacterium carotovorum* (positive control) and Metchnikowia *reukaufii* (negative control) on PGA plates, then mixed one colony of each with prepared fresh *E. californica* pollen in BK germination media, in PCR tubes, to provide initial exposure. As a control for polygalacturonase activity from pollen germination itself, we included another tube into which we added a crushed stigma of *E. californica*, as well as a tube with only pollen in BK media. After one hour of co-incubation at room temperature, we checked each treatment for germination and spotted each of these treatments onto fresh PGA plates in triplicate. We also spotted, in triplicate, the microbes themselves directly from the PGA plates that they had been grown on. After incubation for 72h days at 37C, CTAB solution (1%) was added for 20m to visualize clearance zones (Fig S2).

### Statistical Analysis

All statistical analyses were done in RStudio 1.2.1335 (61) using R 3.6.2 (62). To compare if pollen germination and bursting differed among microbes, Kruskal-Wallis tests were run (base R) with microbe treatment as predictor of pollen germination or bursting, respectively. Separate tests were performed for each timepoint. To test for differences in pollen germination (and separately, pollen bursting) between microbes, pairwise comparisons between microbes were calculated using the “FSA” package (63) to run a Dunn test with Benjamin-Hochberg p-value adjustment (28 tests). For germination and bursting comparison across dilutions of *A. pollinis* SCC477 (density dependence), one-way ANOVA was calculated at each timepoint and “emmeans” package (64) was used for pairwise comparisons and calculation of p-values, with Tukey adjustment (21 tests).

For comparison of microbial growth in pollen/no pollen treatments, we performed one-way ANOVA at the 24h timepoint, and pairwise comparisons between treatments were calculated with Tukey p-value adjustment (5 tests) (64). Two-way ANOVA was also run, using the initial and final cell counts and pollen treatment as predictors of cell count, followed by pairwise comparisons and p-value calculation with Tukey adjustment (45 tests) to determine whether cell count was significantly changed (increased) over 24h within each treatment (64).

