## Supplementary figures for "Nectar bacteria stimulate pollen germination and bursting to enhance their fitness"

Supplementary Material
Christensen 2021

**Tube lengths of dose dependence assay**

During the dose dependence assay described in the main manuscript, we noticed that the length of the pollen tubes seemed to differ over the treatments. To quantify this observed difference, we went through each of the images at the 24-hour timepoint and measured (by hand in FIJI with the "measure" tool, see supplemental methods 1) every pollen tube in the frame. From these data, we found that the tube lengths of the more dilute treatments (especially 1:20 and 1:100) had pollen tubes that were longer than those in more concentrated (more *A. pollinis* SCC477) treatments (1, 1:2 especially). The control of just sterile BK media had less overall germination (as shown in main manuscript figure 4) and also had very short pollen tubes.

We suspect that this is due to a difference between the triggering of the germination and bursting phenotypes. This difference could be in the concentration of the inducing agent needed to trigger germination vs bursting or that it is due to an entirely different, possibly quorum sensing controlled agent that triggers bursting. If a different concentration was needed for germination (low concentration, low threshold) and bursting (high concentration, higher threshold), then we would expect that the high concentration of SCC477 might quickly induce germination and immediately, or soon after, bursting, because it either already had already reached the high threshold of [inducing agent] or did so very quickly after inducing germination. With continued increasing dilutions, it may take longer for induction of germination, and longer still for the [inducing agent] to reach levels that induce bursting of the tubes, allowing the tubes to grow to longer and longer lengths before they are burst and halted in their growth. This is also what we see in Fig 4 of the main manuscript, with the bursting rate lagging behind the germination rate for each treatment, especially at the lowest [SCC477] treatments.

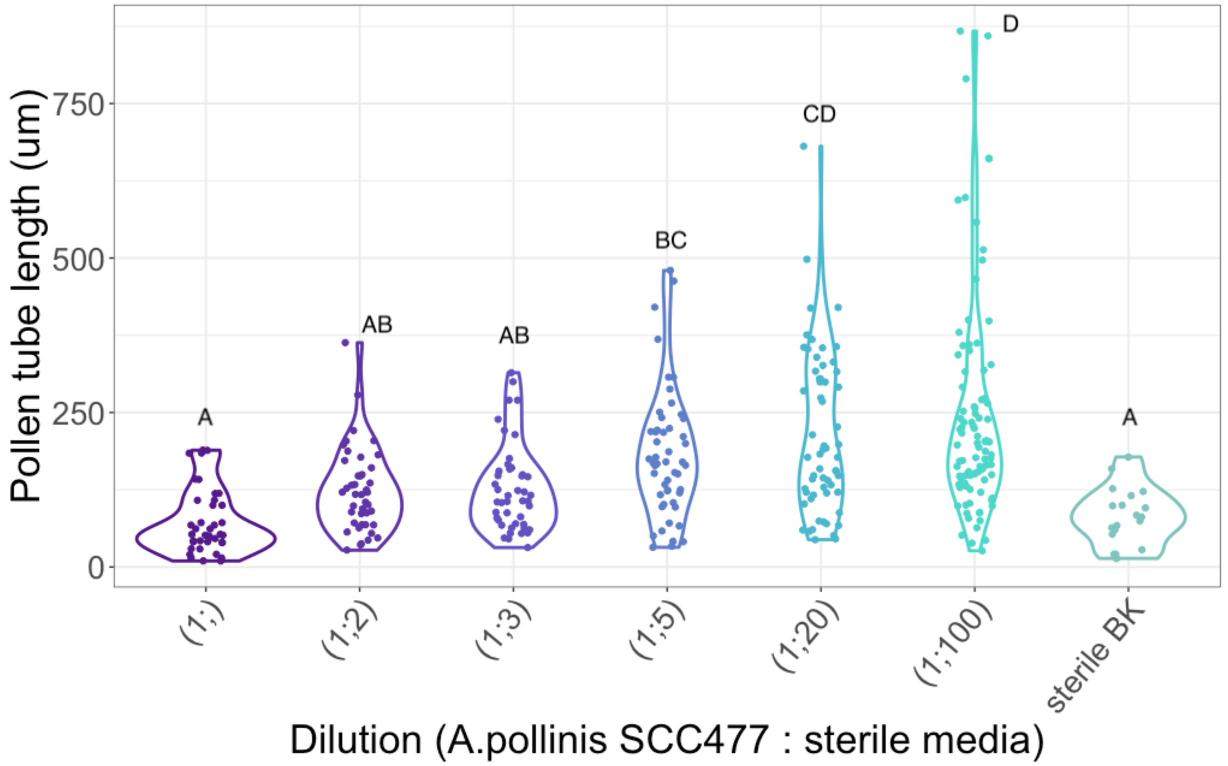


**Figure S1: Pollen tube lengths at 24h are dose dependent.** All pollen tube lengths were measured in every image of the dose dependence assay; the 24h timepoint measurements are shown here. Treatment labels correspond to SCC477 dilution in sterile BK media. Letters indicate significance p<0.05 with ANOVA, multiple comparisons with Tukey p value adjust. Each treatment includes three biological replicates (wells). Average 55.2 pollen tubes measured per treatment.

**Polygalacturonase activity assay**

We initially hypothesized that the polygalacturonases found in the genome of SCC477 may play a role in pollen germination or bursting phenotype. To test whether polygalacturonases found in the *A. pollinis* SCC477 genome are active, and whether they have (qualitatively) more activity upon exposure of the strain to pollen, we utilized a plate-based PGA assay. We grew SCC477, *Pectobacterium carotovorum* (positive control) and *Metchnikowia reukaufii* (negative control) on PGA plates (.1M Tris-HCl, 1.5% agar, 0.5% yeast extract, and 0.5% PGA), then mixed one colony of each with prepared fresh *E. californica* pollen in BK germination media, in PCR tubes, to provide initial exposure. As a control for polygalacturonase activity from pollen germination itself (plant derived pectinases), we included another tube into which we added a crushed stigma of *E. californica* (which induces germination), as well as a tube with only pollen in BK media. After one hour of co-incubation at room temperature, we checked each treatment for germination (y/n) and spotted each of these treatments onto fresh PGA plates in triplicate. We also spotted, in triplicate, the microbes themselves directly from the PGA plates they had been grown on (no pollen). After 3 days at 37°C, Cetyltrimethylammonium bromide (CTAB) (Chem-Impex cat. no 01781) solution (1%) was carefully dripped over the plates, specifically around the colonies. We waited 20 minutes and then checked for clearance zones. CTAB stains PGA milky white, so clearance zones (unstained) indicate that the spotted colony created and released polygalacturonases into the media. There did not appear to be any difference with addition of pollen in any treatment, and only the positive control spots, *P. carotovorum* showed any clearance zones, both with and without pollen.

| | *A. pollinis SCC477* | *P. carotovorum* (+) | *M. reukaufii* (-) | Crushed stigma | BK media only |
|---|---|---|---|---|---|
| + pollen | No clearance (Y) | Clearance (N) | No clearance (N) | No clearance (Y) | No clearance (N) |
| - pollen | No clearance | Clearance | No clearance | X | X |

+ pollen

- pollen

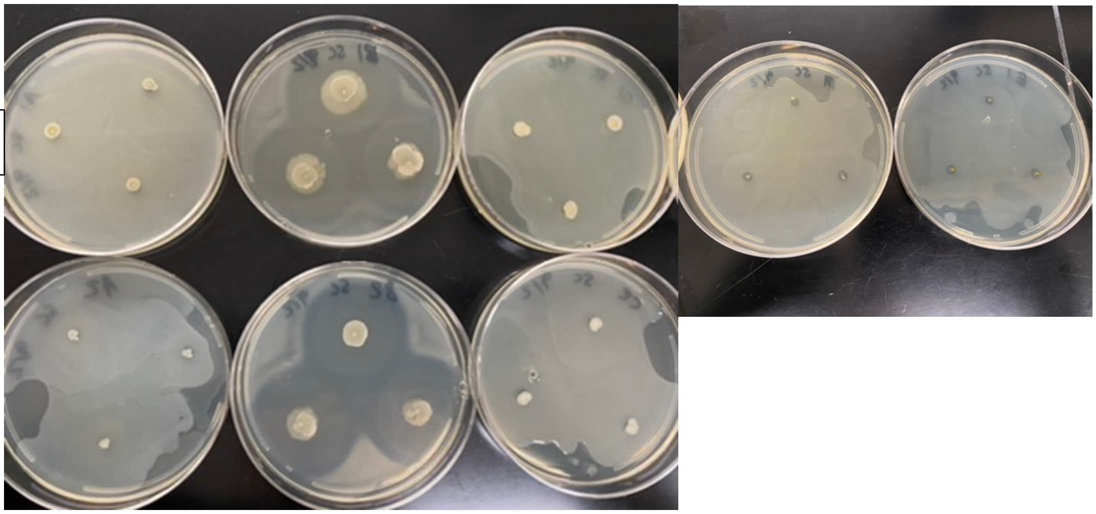

**Figure S2: PGA activity not linked to germination phenotype**. 'Clearance' or 'No clearance' indicates polygalacturonase activity and undetectable activity, respectively. (Y)/(N) in "+ pollen" row indicates whether the pollen was observed to be germinating (Y) or not (N) after 1h in that treatment, just before spotting onto PGA. The images below the table are in the same order/orientation as in the table. The faint shadows around the spots in the 'crushed stigma' and 'BK only' plates are a result of marking the placement of the spots with sharpie on the other side of the plate, not clearing of CTAB.

### PGA degradation

We also wanted to test the polygalacturonase activity of strains used in the pollen germination assays to qualitatively determine whether the differences in induction phenotype were a result of differing polygalacturonase activity. For these tests, we grew each strain on R2A plates (or YM for *M. reukaufii)* then, as above, spotted them onto PGA plates, grew them at 37°C for 2-3 days, and gently dripped CTAB (1%) around the colonies, then waited 20 minutes to check for clearance zones.

Our results are summarized below (Table S1). As expected, our positive control, *P. carotovorum,* had distinct clearance zones in all replicates, and our negative control, *M. reukaufii,* had none. The *Acinetobacter* strains showed mixed results, with SCC477 and SCC474 showing slight clearance zones on one plate (3 spots) but not on the other two plates (6 spots). Given the above results (Figure S2), and the lack of any discernable activity for *A. boissieri*, which also had high induction, we do not believe that this slight activity in some replicates plays a role in germination induction phenotype.

| Strain: | Species: | Clearance zone: | Total reps: | Reps showing clearance: | Accession numbers and locus tags of pectinase genes | Notes: |
|---|---|---|---|---|---|---|
| *P. carotovorum* | *P. carotovorum* | Clearance | 9 | 9 | | Positive control |
| EC52 | *M. reukaufii* | No clearance | 9 | 0 | | Negative control |
| SCC474 | *A. pollinis* | Slight | 9 | 3 | | |
| SCC477 | *A. pollinis* | Slight | 9 | 3 | | |
| ANC4422 | *A. boissieri* | No clearance | 9 | 0 | NZ_FMYL01000001.1, BLS38_RS01625 BLS38_RS12295 | |
| BB362 B1 | *A. nectaris* | No clearance | 9 | 0 | NZ_KI530734.1, P256_RS09450; NZ_KI530712.1, P256_RS02650 | |
| *A. apis* | *A. apis* | ND | | | NZ_FZLN01000001.1 SAMN05444584_0345, NZ_FZLN01000002.1 CFY84_RS07245 | |

**Table S1: Polygalacturonase activity of strains used in this manuscript.**
