## Supplementary methods 1 for "Nectar bacteria stimulate pollen germination and bursting to enhance their fitness"

### Supplementary methods 1: counting and classifying pollen grains

##### Macro script for counting total pollen grains ~40um diameter in Fiji2 (v2.0.0)

We validated this macro script to be accurate with a paired t-test comparing macro script counts with hand counts of the same images (90-minute timepoint from three assays). The macro counts and hand counts were not significantly different (N=53, p= 0.3353). The 90 minute timepoint was selected because the pollen tubes and burst grains present at 90m (in some images) were initially an obstacle for accurate macro counting.

All images for each timepoint were analyzed in a batch using “Process”-> “Batch”-> ”Macro” with the following script:

```
run("8-bit");
setAutoThreshold("Default dark no-reset");
//run("Threshold...");
//setThreshold(176, 255);
run("Make Binary", "thresholded remaining black");
run("Convert to Mask");
run("Fill Holes");
run("Watershed");
run("Analyze Particles...", "size=900-3500 circularity=0.30-1.00 show=Overlay
display exclude summarize");
```

Note: The size and circularity ranges can be changed for use with different pollen sizes and shapes. This is especially important in the later time points (with germination) as the circularity measure can be altered by protruding pollen tubes, or even bulging pollen tube tips.

##### Classification of pollen

After counting total pollen grains, the Plugin “CellCounter” (https://imagej.nih.gov/ij/plugins/cell-counter.html) was used to count pollen grains within each of three classifications (Fig 1). Any pollen grain touching the edge of the photo was excluded in both the total and subset counts. Grains were counted as “germinated” if any pollen tube was visible and not burst, shown here as the green-labeled grains “1”. If protoplasm was spilling out of a pollen grain without a visible pollen tube, they were counted as “burst” “2”; if protoplasm was visibly exiting the pollen tube tip of a germinated grain, they were instead counted as “tip burst”, labeled in orange “3”.

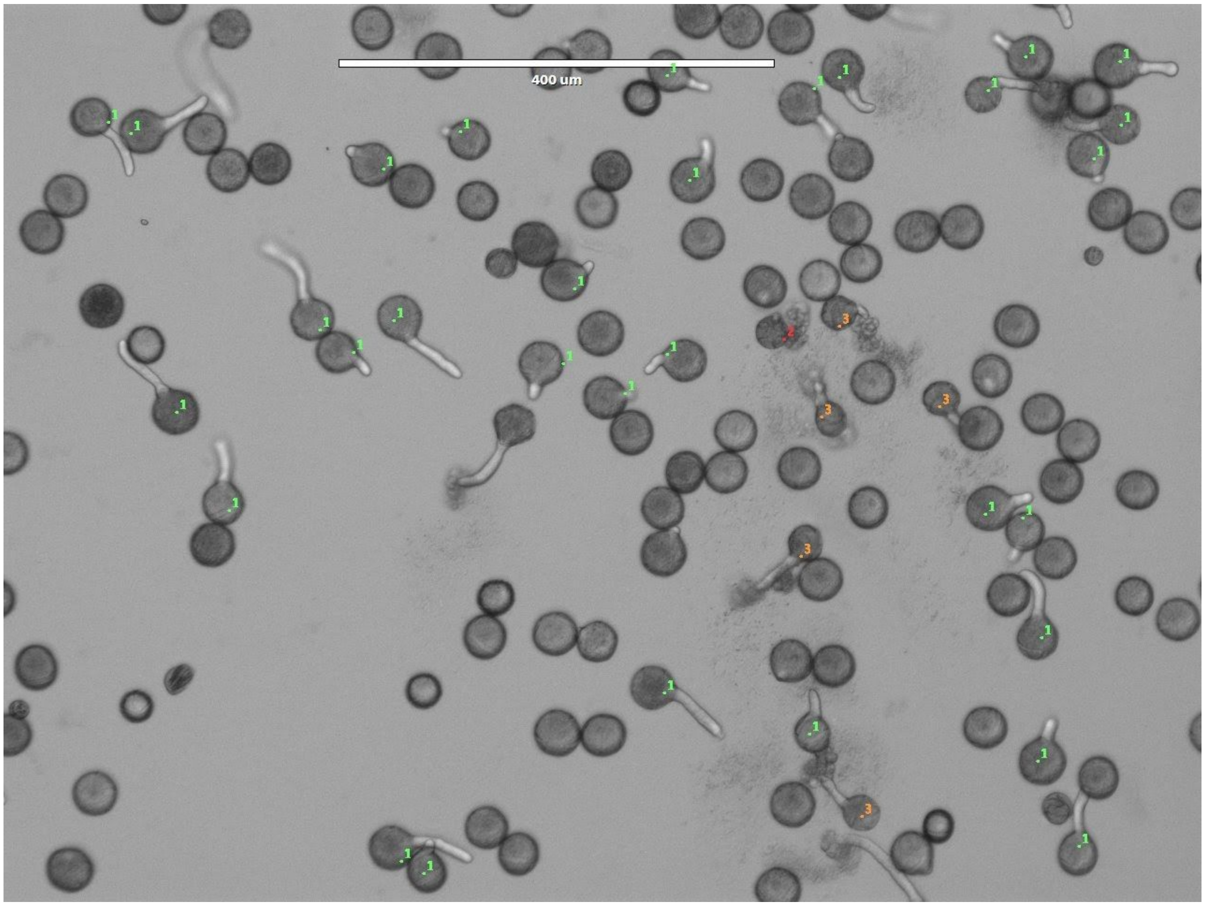


**Measuring pollen tube lengths (dilution/dose-dependence only)**
Using the basic FIJI tool, we initially use the image scale bar to set the scale, then use ROI manager (“Analyze”->“Tools”->”ROI Manager”) with Freehand drawing tool to trace and keep track of pollen tubes (check “Show all”) “add [t]” each line to the manager, then highlight all and “Measure”.

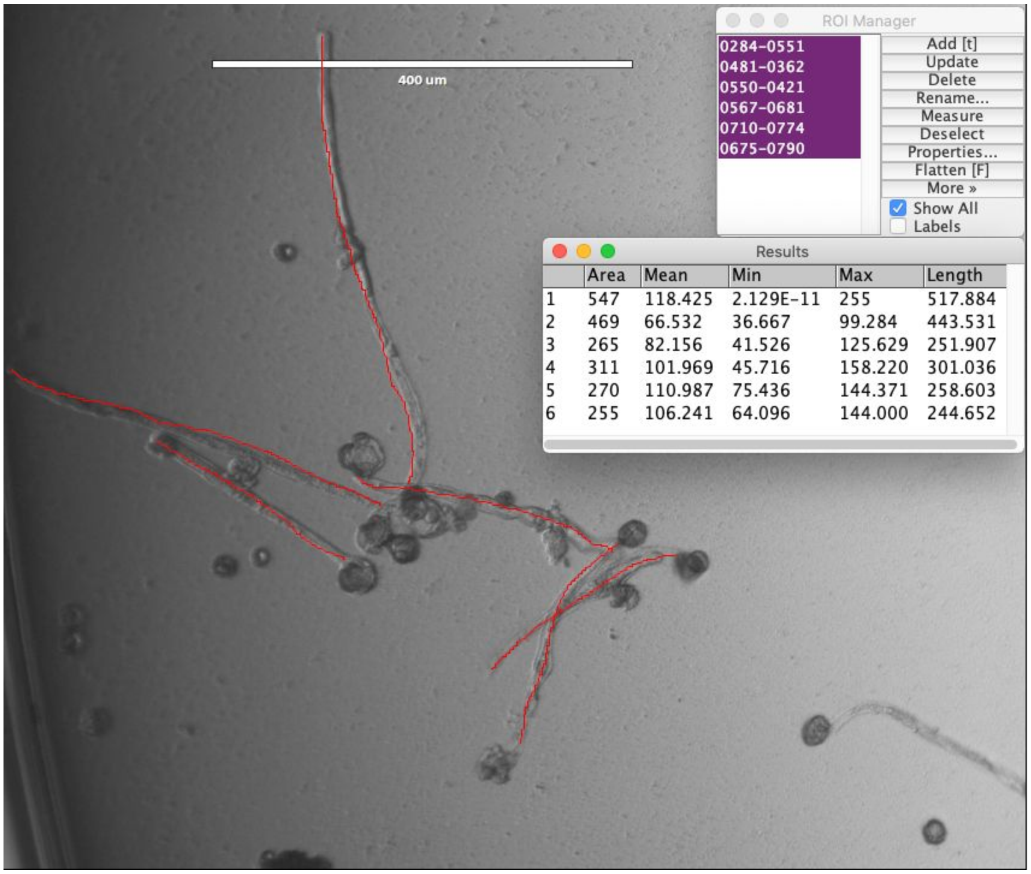
400 um
ROI Manager
0284-0551
0481-0362
0550-0421
0567-0681
0710-0774
0675-0790
Add [t]
Update
Delete
Rename...
Measure
Deselect
Properties...
Flatten [F]
More »
Show All
Labels
Results
Area Mean Min Max Length
1 547 118.425 2.129E-11 255 517.884
2 469 66.532 36.667 99.284 443.531
3 265 82.156 41.526 125.629 251.907
4 311 101.969 45.716 158.220 301.036
5 270 110.987 75.436 144.371 258.603
6 255 106.241 64.096 144.000 244.652
