## Supplementary methods 2 for "Nectar bacteria stimulate pollen germination and bursting to enhance their fitness"

**Supplementary methods S2**

*Microbe identification*

Within 1 day after select assays (2x microbial impact on pollen assays , 1x dose dependence assay), used wells (50ul) were plated onto separate plates of Tryptic Soy Agar with cycloheximide and allowed to grow for 48-72 hours. For each morphotype present, (if more than one) material from one or two representative colonies was selected and spotted in duplicate onto an MTP 384 Ground Steel Target Plate (Bruker Daltonics), overlaid with 1uL 70% formic acid, and allowed to air dry. 1uL of alpha-cyano-4-hydroxycinnamic acid (HCCA) matrix dissolved in a 50:45:5 solution of acetonitrile, water, and trifluoroacetic acid was added and let dry [same prep as in Seuylemezian 2018, JPL microbes paper. doi: 10.3389/fmicb.2018.00780]. Spectra were obtained twice per spot, in succession, using an ultrafleXtreme MALDI-TOF instrument (Bruker Daltonics, Billerica, MA, United States). Spectra for each isolate were then compared with a custom in-house library of main spectral profiles (MSPs) and the Bruker libraries (Bacteria, Eukaryotes) using MBT Compass Explorer software (Bruker Daltonics, Billerica, MA, United States). Log scores between 1 and 3 were generated for each isolate's top 10 best matches. Scores ≥2.0 are considered close matches; lower scores indicate that the exact species may not have been present or matched closely enough to the library spectra. Genus identification was used for those with scores above 1.6. Sometimes, there is no spectra generated from a spot (HCCA crystals do not form correctly, the spotting is too thick, the laser strikes areas where either of these causes lack of ionization), and this leads to lack of identification for the microbes in those wells. Spots/wells that failed to generate spectra were removed from the compiled 'MALDI_identifications.csv'.

*Overall MALDI results*

All identified *Acinetobacter* had scores >1.6. Of the 64 spectra generated from *Acinetobacter* inoculated wells, 82% returned *Acinetobacter* (genus), and of those, 85% were also accurate to species inoculated. All morphologically distinct colonies were tested, so the 64 generated spectra include some wells for which both *Acinetobacter* and another contaminant were identified (see MALDI_identifications.csv)

Six of the seven spectra from wells inoculated with *Pectobacterium carotovorum* were identified as *P. carotovorum*, scores approximately 1.6, just at or below genus-level confidence. We did not find *Pectobacterium* as a contaminant in uninoculated controls or *Acinetobacter*-inoculated controls.

In one assay, one of the plates from a control well was contaminated with *A. pollinis SCC477*. It was the only colony on the plate, indicating that contamination may have happened very late in, or after, the assay or during plating. This well did not exhibit greater germination/bursting than the other wells by 90m. The same assay showed the same contamination of one well of *M. reukaufii* with *A. pollinis SCC477,* and also no effect on pollen, indicating that the contamination may have happened after the assay, or in an amount small enough to not produce the phenotype that was seen in other wells.

*M. reukaufii* was identified visually in the images of the final timepoint, as it is easily seen, distinctive, and was never seen in control pollen or any other wells. It was present in all the wells it was added to. The wells were still plated on TSA (with cycloheximide) to identify any bacterial contamination.

Overall, contaminants identified with support to the genus level (score between 1.6 and 2) were *Rosenbergiella* and *Filifactor.* Contaminants identified **without strong support** (score <1.6) were *Agromyces, Rahnella, Cupriavidus, Ewignella, Halomonas, Citrobacter, Neisseria, Thauera,* and *Pantoea*. All contaminants only appeared once, aside from *Agromyces* (score <1.6) and *Rosenbergiella* (score between 1.6 and 2).

*MALDI results by assay*

In the dose dependence assay (1.16.20), the treatments with doses of 1, 1:2, 1:3 and 1:20 all re-identified the inoculated *A. pollinis* SCC477 with scores between 1.87-2.54. Treatment 1 (full dose of SCC477) also returned the contaminant *Agromyces* (low score, 1.23) in one well. The 1:5

treatment had only one spot that resulted in usable peaks (of 6 total spots/treatment), which was matched to *Citrobacter* with a low score (1.2), though very sticky, characteristically *Acinetobacter pollinis*-like colonies were also obtained, indicating problems with the matrix or spotting thickness. The same was true for the 1:20 treatment spots, which returned three low-score contaminants (*Ewignella, Halomonas, Cupriavidus*, scores 1.22-1.29) but also *A. pollinis*, with a score of 1.98 (close to species level confidence). The 1:100 treatment had only one spot that had usable spectra, which was identified (with very low scores 1.08, 1.12) as first *Thauera* and then *Agromyces*. But again, all plates from each well had the majority of colonies of the very sticky and characteristic colony morphology of SCC477.

In the 10.25.19 microbial impacts on pollen assay, two wells of each treatment were plated, and the colonies identified with MALDI_TOF. *A. pollinis* SCC477 was correctly identified from the treatment to which it was added (scores 2.32-2.64). The *A. apis* treatment had 7 of 8 spectra (4 spots, 2 spectra per spot) identified as *A. apis* (1.96-2.52), and one as *A. nectaris* (1.89). The *M. reukaufii* treatment, as described previously, was visually confirmed to contain the yeast (readily seen under microscope, not present in any other wells) but was still plated on fungicide-containing TSA to look for bacterial contamination. The plate from one well had no bacterial growth, and the second had very few colonies, but were identified as *A. pollinis* SCC477 (scores 2.22-2.61) indicating that there may have been some late cross-contamination of that well, or that SCC477 was present on the pollen in very low numbers from the beginning. 7 of 8 MALDI passes (4 spots) generated spectra from the *P. carotovorum* plates, which were all identified as *Pectobacterium,* albeit with scores of only 1.44-1.77 (low scores, one to genus level). As discussed above, the control well had only one colony grow on the plate, and this was identified as *A. pollinis* SCC477. The other well had no colonies grow. The control well and *M. reukaufii* well that had (low) SCC477 contamination were in the same column of the plate.

For the 8.15.19 microbial impacts on pollen assay, every well was plated and spotted once, 2 spectra read per spot. For *A. pollinis* SCC477 inoculated wells, 3 of 4 wells had colonies that generated spectra, and all spectra gathered matched *Acinetobacter* with genus-level support

(1.86-2.06), one, matching SCC 477 had species level support. *A. apis* inoculated wells generated 4 spectra from 2 wells, all of which matched to *Acinetobacter* with at least genus level support (1.69-2.16). *A. nectaris* inoculated wells all generated two spectra (8 total) with 6 of these matching *A. nectaris* with species-level support(2.02-2.3), one with genus level support (1.79) and one matching SCC477 (2.13). The controls and one of the *M. reukaufii* wells all showed some contamination, which was identified with low to genus level support (1.5-1.77) as *Rosenbergiella* (7 of 8 spectra) indicating that this was already present on the pollen. *Providencia* was identified with very low (1.43) support from one spectra.

More details and exact scores in 'MALDI_identifications.csv'
